# Iron regulatory pathways differentially expressed during *Madurella mycetomatis* grain development in *Galleria mellonella*

**DOI:** 10.1101/2022.12.20.520897

**Authors:** Imad Abugessaisa, Mickey Konings, Ri-Ichiroh Manabe, Tsugumi Kawashima, Akira Hasegawa, Chitose Takahashi, Michihira Tagami, Yasushi Okazaki, Wilson Lim, Annelies Verbon, Ahmed H. Fahal, Takeya Kasukawa, Wendy W.J. van de Sande

## Abstract

**Background:** Mycetoma is a neglected, chronic granulomatous infection of the subcutaneous tissue, most often caused by the fungal pathogen *Madurella mycetomatis*. Characteristic of the infection is the formation of grains. However, knowledge of the function and formation of the grain is limited. To map the processes leading to *M. mycetomatis* grain formation, we used a *Galleria mellonella* larvae infection model and time-course transcriptomic profiling.

**Methods:** *Galleria mellonella* larvae were infected with *M. mycetomatis* genome strain mm55. At 4h, 24h, 72h and 168h post-inoculation, RNA was extracted from larval content. Two types of sequencing libraries were prepared for time-course transcriptomic profiling and analysis.

**Findings:** In the infected *G. mellonella*, 88.0% of the RNA sequence reads mapped to *G. mellonella*, while only 0.01% mapped to *M. mycetomatis*. Differential Gene Expression analysis revealed that 3,498 *G. mellonella* and 136 *M. mycetomatis* genes were differentially expressed during infection. Most of the enriched GO terms of both host and pathogen are linked to energy pathways, nucleobase metabolic process as well as cation and iron transport. Genes related to iron transport were highly expressed by both *G. mellonell*a (transferrin and ferritin) and *M. mycetomatis* (SidA, SidD and SidI). A protein-protein interaction network analysis of *D. melanogaster* homologous genes in *M. mycetomatis* revealed the expression of the entire siderophore biosynthesis pathway throughout infection.

**Interpretation:** The identification of the importance of iron acquisition during grain formation can be exploited as a potential novel diagnostic and therapeutic strategy for mycetoma.

**Research in context:** *Evidence before this study:* Mycetoma is a chronic, neglected tropical infectious disease, characterised by a large subcutaneous mass and the formation of black grains in the affected tissue. Treatment for mycetoma is disappointing as in 25-50% of the patients recurrences are noted and up to 15% of patients will have to undergo amputation. The main reason behind this poor treatment response is the formation of protective structures by the pathogen upon entering the human body. These structures are called grains and provide a strong barrier for antifungal agents. Although grains are the hallmark of mycetoma, it is currently not known how these grains are formed. To improve the current therapy, it is important to gain insights in grain formation.

*Added value of this study:* We unravel the processes leading to grain formation and development in an invertebrate model of *Madurella mycetomatis* grain. We were able to build a model of grain formation and demonstrated that iron sequestering plays an important role in this process. Our findings were an important milestone in understanding the pathogenesis of mycetoma which has been a mystery for decades.

*Implications of all the available evidence:* The findings, will provide leads for future drug development of mycetoma treatment and therefore, improve patients live and end the need for amputations.

## Introduction

Mycetoma is a neglected disease endemic in tropical and subtropical regions. It is characterized by the formation of subcutaneous tumour-like swellings and grains that spread to deep tissues and skin. The extremities are affected most, but no body part is exempted. (**Figure 1A)** (1, 2). Grains are biofilm-like structures in which the causative agent resides **(Figure 1B)** (3). The disease progression is gradual, and symptoms develop over the course of months to years. Although mycetoma can be caused by several causative agents, in more than 40% of all published cases, the fungus *Madurella mycetomatis* was the causative agent (4, 5).

**Figure 1:**
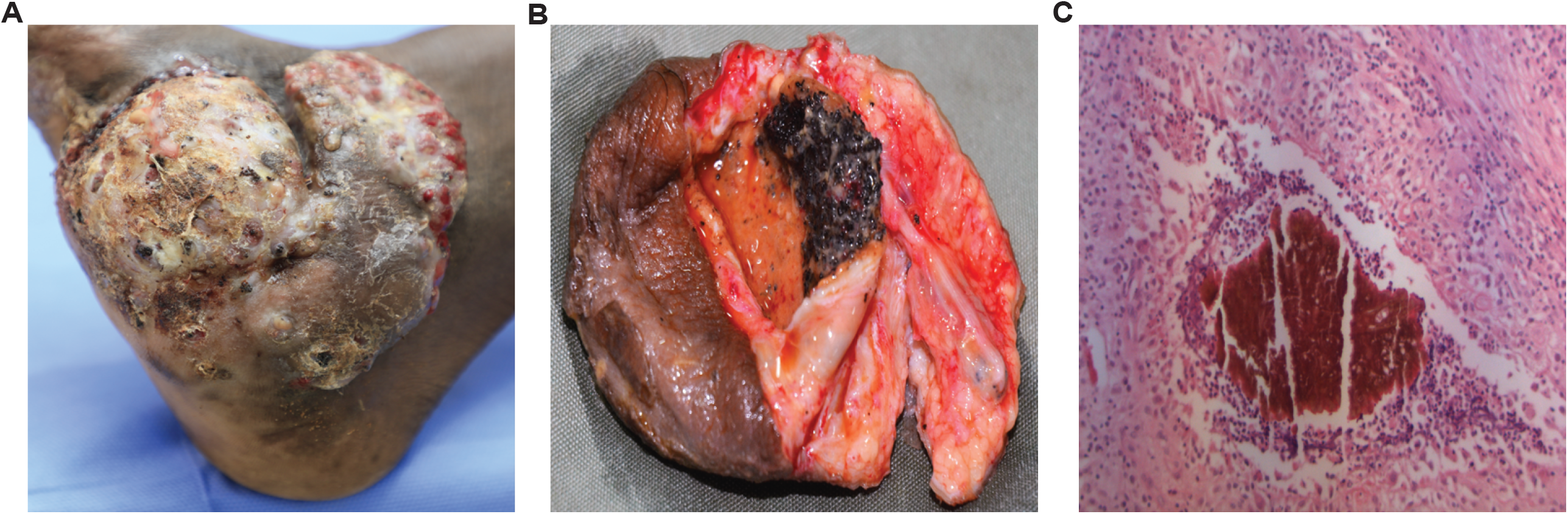
Mycetoma of the foot with characteristic mycetoma grains. **A**. An extensive case of mycetoma on the foot. **B**. Black grains of *M. mycetomatis* are clearly visible inside the infected tissue. **C**. A histopathological picture of the *M. mycetomatis* black grain, with *M. mycetomatis* hyphae embedded in cement material (H&E staining, 40 times magnified using H&E staining).

Although mycetoma grain formation is one of the hallmarks of mycetoma, currently there is little known about their function and formation. Some studies demonstrated that grain components could protect the causative organism against the host immune response and antifungal therapy (6, 7). Others have demonstrated that these grains consist of a cement-like material, mainly composed of heavy metals, melanin, lipids and proteins (3).

Grains can only be formed in *in vivo* models. The simplest model in which grain formation can be studied is the *M. mycetomatis* grain model developed in larvae of the invertebrate *Galleria mellonella* (8, 9, 10). *G. mellonella* has been widely accepted as a model organism as the larvae are easy to use, inexpensive, have an innate immune system which is comparable to that of mammals, are free of the legal/ethical restrictions associated with the use of mammals, and most importantly the obtained results closely correlate with those obtained in mice (11, 12, 13). In *G. mellonella, M. mycetomatis* grains can be formed which resemble the grains in mouse models and human patients (14). In a previous study using this model, we demonstrated that this model can be used to follow grain formation over time and that differences in protein abundance could be demonstrated by using a label-free quantitative LC-MS/MS proteomic approach (15). However, which transcriptional responses are important in the process of grain formation are currently unknown.

In this study, we aimed to profile the transcriptomic changes of both the larval host and the *M. mycetomatis* pathogen during grain formation to enhance our understanding of grain formation. We found differentially expressed pathways in both the larval host and *M. mycetomatis*, among which we identified the interplay in iron regulation between host and pathogen.

## Material and methods

### Culturing and inoculum preparation of M. mycetomatis strain MM55

*M. mycetomatis* genome strain MM55 was cultured on Sabouraud Dextrose Agar for two to three weeks at 37□C. Mycelium was harvested, sonicated at 28 micron (Soniprep 150, Beun de Ronde, the Netherlands) and incubated at 37□C in RPMI-media supplemented with 0.35 g/L L-glutamine, 1.98 mM 4-morpholinepropanesulfonic acid (MOPS) and 100 mg/L chloramphenicol. After two weeks, the mycelia were separated and washed by vacuum filtration (Nalgene, Abcoude, the Netherlands using a 22 micron filter (Whatman)). The mycelial biomass was scraped from the filter, the wet weight was determined, and a suspension was prepared containing 100 mg mycelial biomass per ml phosphate-buffered saline (PBS). The suspension was sonicated at 28 micron for 2 minutes (Soniprep 150, Beun de Ronde, the Netherlands), the resulting homogeneous suspension was washed once in PBS and again diluted to a concentration of 4 mg per 40 μL PBS, corresponding to 600-850 CFU/larvae.

### Infection of G. mellonella larvae with M. mycetomatis and RNA isolation

*G. mellonella* larvae were obtained from Terra Equipment Voedseldieren (Cuijk, The Netherlands) and kept in the dark on wood shaving at room temperature until use. Within five days of receipt, larvae of approximately 300-500mg were selected for experimental use. The selected larvae were divided over Petri dishes containing 90mm Whatman filter paper and five larvae per dish. 40 μL of the prepared inoculum of *M. mycetomatis* strain MM55 was injected in the last left proleg of the larvae using an insulin 29G U-100 needle (BD diagnostics, Sparsk, USA), resulting in a final concentration of 4 mg fungal biomass/mL. To monitor the course of the infection, a separate group consisting of 15 larvae was infected, and survival was recorded daily for the duration of ten days. During all experiments, Pupa were removed from the equation, and non-infected larvae were included as control. At 4, 24, 72 and 168 hours post-infection, the contents of three larvae were pooled and flash-frozen with liquid nitrogen, followed by mechanical crushing using a pestle and mortar. The resulting powder was suspended in RLT buffer (supplemented with 1% β-Mercaptoethanol), provided in the RNeasy Mini Kit (Qiagen, Germany), and incubated at 57□C for 3 minutes. RNA isolation was further continued according to the manufacturer’s instructions. The presence and quality of RNA were assessed by NanoDrop and gel-electrophoresis.

### Determination of the fungal burden in G. mellonella

At 4, 24, 72 and 168 hours post-infection, larvae were sacrificed, and haemolymph was harvested. Melanisation of haemolymph was determined by diluting the haemolymph 1:1 in (IPS). The optical density was measured at 405 nm using the Nanodrop (Thermo Scientific, USA). To fixate the larvae, 100 μL 10% buffered formalin was injected into the larvae. The larvae were moved into 15 mL Greiner tubes containing 5 mL 10% buffered formalin. After 24 hours of fixation, whole larvae were longitudinally dissected into two halves using a scalpel and fixated in 10% buffered formalin for another 48 hours before further routine histological processing (16). The two halves were stained with hematoxylin and eosin (HE) and Grocott methenamine silver for further histological examination. The fungal burden analysis detailed in in (**Supporting Information Text**).

### RNA sequencing and LQ-ssCAGE library preparation, sequencing, mapping, and processing

As illustrated in **Figure 2A**, we prepared two types of libraries for high throughput profiling, RNA-Seq and Low Quantity single strand CAGE library preparation (LQ-ssCAGE) libraries. Full list of samples is shown in **Supplementary Table S1**. RNA-Seq libraries were prepared using 1 ug of total RNA with TruSeq Stranded mRNA Library Prep kit (Illumina) following the manufacturer’s instructions. LQ-ssCAGE sequencing libraries were prepared following the procedures described in (17). Sequencing specifications of RNA-seq and LQ-ssCAGE libraries and mapping and quantification of the raw sequence reads detailed in (**Supporting Information Text**). Using LQ-ssCAGE, a genome-wide map of Transcription start site (TSS) was established for the *Galleria mellonella* larvae and *M. mycetomatis* by precisely identifying promoters and enhancers as described in (**Supporting Information Text**).

**Figure 2:**
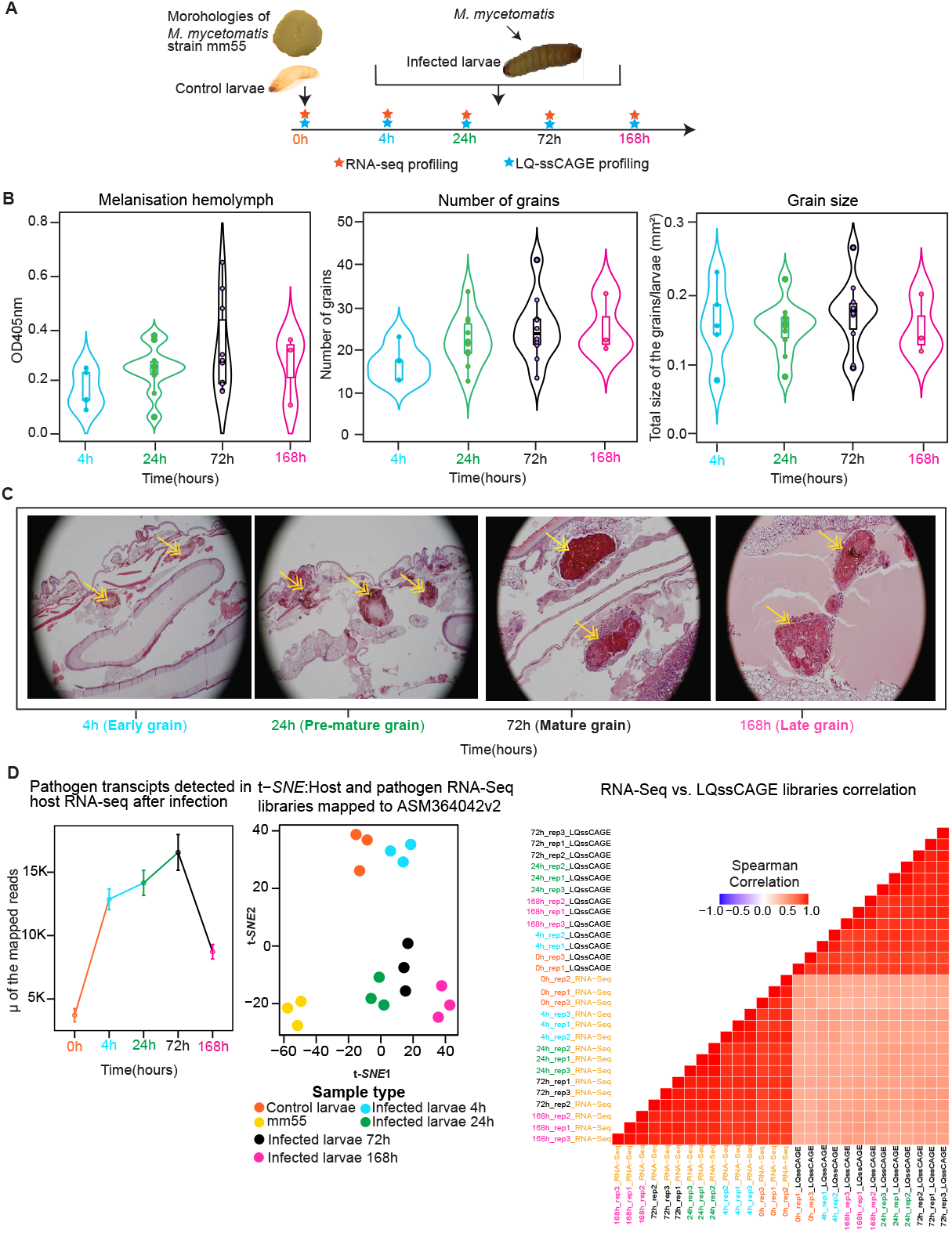
Experimental workflow, burden of infection on *G. mellonella* larvae, and quality of transcriptomics data. **A**. Schematic illustration of the time course experiment design and transcriptomic assays used for high throughput RNA profiling. Total RNA was extracted at five-time points (three replicates each). At time point 0, RNA was extracted from healthy *G. mellonella* larvae (host) and cultured *M. mycetomatis* strain (mm55) (pathogen) to serve as non-infected controls. After the inoculation of *G. mellonella* larvae with *M. mycetomatis*, RNA was extracted at 4h, 24h, 72h and 168h post-infection. **B**. To evaluate the burden of infection in the host, the melanisation of the hemolymph, the number of grains and the grains size were measured and plotted in violin plots for each time point. The violins are colored magenta for t=4h, green for t=24h, black for t=72h and pink for t=168h. The highest values are noted at t=72h (the maximum burden of infection). **C**. Grain formation in the host was visualized 40 times magnified using H&E staining and light microscopy. Yellow arrows point towards the grain inside the capsule. Based on the characteristics of the grain development observed at each time point, the grains were defined as early, pre-mature, mature, and late grains. In the early grains, the cement material is not formed, and hemocytes are present between hyphae. In the pre-mature grains, hemolymph is forming, but still, individual hemocytes can be noted within the forming cement material. In the mature grain the cement material is completely formed and a capsule surrounds the grain. In the late grain, you see the capsule disappearing and an influx of hemocytes towards the grain. **D**. Quality of the RNA sequencing and LQ-ssCAGE dataset. The line plot shows the number of the pathogen reads detected in the host RNA-seq data after infection, the largest number of the pathogen transcripts detected at time 72h (mature grain) (lines are colored by time point). Data in the line plot are represented as mapped sequnced reads ± SEM.Result of t-*SNE* clustering. The replicates at each time point are shown as colored dots. The t-*SNE* plot shows the clustering of the RNA-seq libraries mapped to the host genome. RNA-Seq vs. LQ-ssCAGE libraries correlation, the correlation matrix shows the Spearman correlation of the RNA-Seq and LQ-ssCAGE reads between replicates.

### Differential expression analysis

Raw RNA-Seq counts were normalized using the edgeR R data package (18). First, we generated the DGElist object and then calculated the normalization factor for the raw read using the calcNormFactors function and trimmed the mean of M value (TMM). The recommended count-per-million (CPM) was obtained as a normalized gene expression matrix. The normalized expression values were used for downstream analysis to identify differentially expressed genes and perform advanced computational analysis of enriched pathways which was detailed in (**Supporting Information Text**).

To process time course data and understand the overtime changes in the eumycetoma grain in the infected larvae, a differential gene expression analysis workflow was designed (**Supplementary Figure S1**). This workflow was used to analyse two types of changes. First, the sharp change analysis, which detects the changes between pre-infection and post-infection (e.g., T0 and T168). The sharp changes were detected using a two steps regression model analysis implemented in maSigPro R package (19). The result of this analysis is depicted in **Figure 3A**. The second type of change was the consecutive change, which was detected by computing the changes between consecutive time-points and between every time point and T0. To detect consecutive changes, a linear model analysis (LIMMA) embedded in edgeR R package was used.

**Figure 3:**
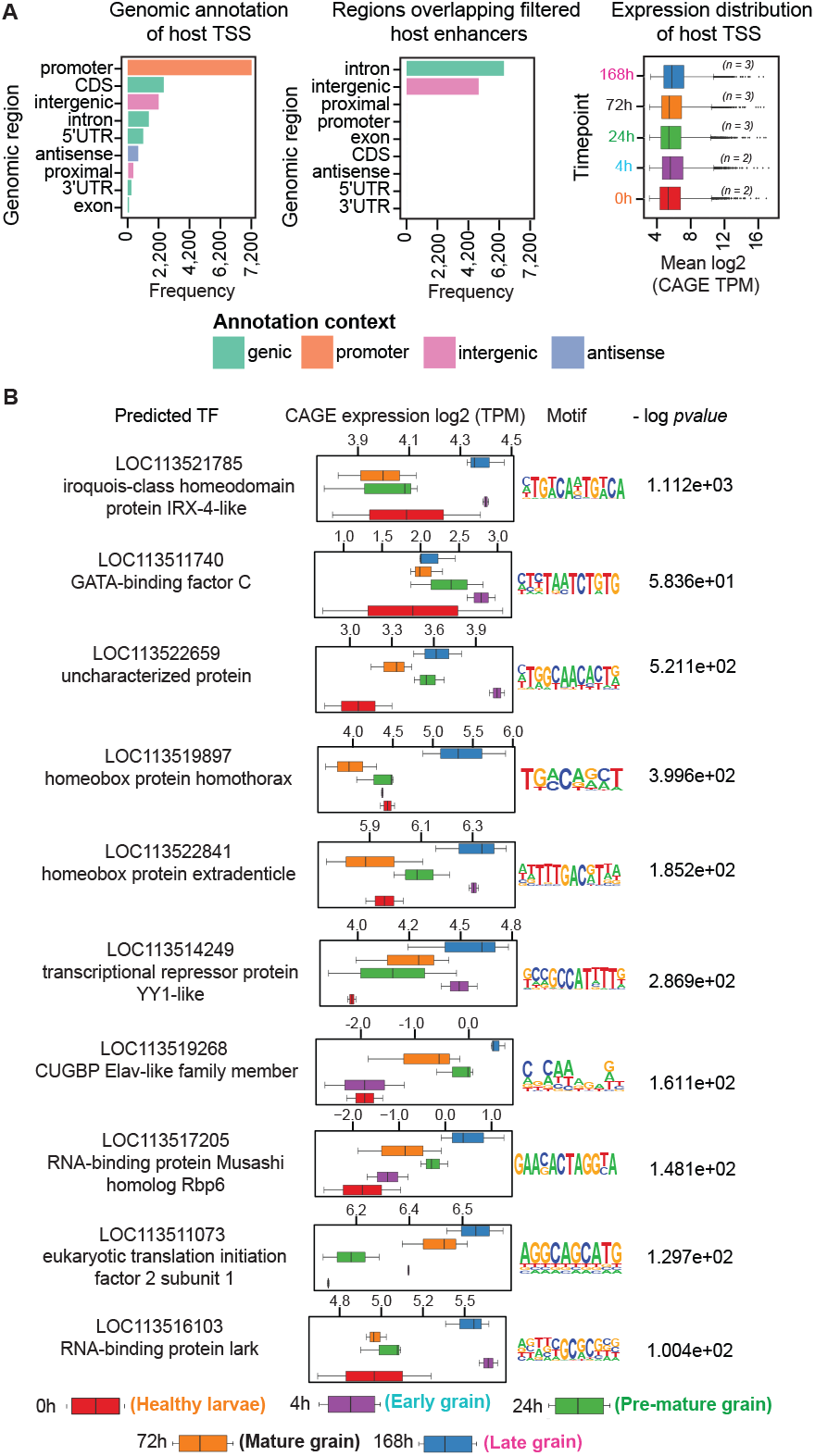
Landscape of the promoters and active enhancers of the host genome and the predicted transcription factors. **A**. Barplots of genomic annotation of the TSS and the regions overlapping filtered enhancers. The bars are colored by annotation context. The majority of the identified TSS peaks are annotated as promoters. Active enhancers are annotated as intronic and intergenic regions. The expression distribution of TSS shows minor differences of expression per time point illustrated. Expression distribution of host TSS represented as boxplot with lower whisker (represents smallest observation greater than or equal to lower hinge in the boxplot - 1.5 * IQR), median and upper whisker (represents largest observation less than or equal to upper hinge in the boxplot + 1.5 * IQR).The largest mean CAGE TPM expression was observed at 168h. **B**. Predicted transcription factors (TF). TF predicted using Homer motif analysis (Material and Methods) for each of the TF CAGE expression and colored by time point. The CAGE expression shown as boxplot with lower whisker (represents smallest observation greater than or equal to lower hinge in the boxplot - 1.5 * IQR), median and upper whisker (represents largest observation less than or equal to upper hinge in the boxplot + 1.5 * IQR).Sequence log for each motif is shown together with the -log *p-value*. Full list of all predicted TF and associated GO terms in (**Supplementary Table S3**).

### Identification and analysis of 1,8-Dihydroxynaphthalene (DHN)-melanin, pyomelanin and siderophore biosynthesis pathways in M. mycetomatis

We investigated the regulation of iron regulation in *G. mellonella* in three biosynthesis pathways in the *M. mycetomatis*. The three pathways are the 1,8-Dihydroxynaphthalene (DHN)-melanin, pyomelanin and siderophore. The detailed methods for the identification and analysis of these pathways in (**Supporting Information Text**).

### Ethics statement

We used the invertebrate *G. Mellonella* larvae this model free of the legal/ethical restrictions associated with the use of mammals

### Statistical Analysis

To assess the burden of infection in the *G. mellonella* we used GraphPad Prism version 8.4.3 (GraphPad Software, LLC). For the rest of the analysis, we used R version 4.1.2 (2021-11-01). Data are represented as either in box-and-whisker plots, line plot or bar plots (represented as expression ± SEM) as specified in the figure legends. For testing differentially expressed genes, F-statistic, the associated P value, and the adj. P value were corrected using Benjamini-Hochberg multiple testing correction.

## Results

### Maximum burden of infection observed three days after inoculation

To dissect the *M. mycetomatis* grain development in *G. mellonella*, we infected *G. mellonella* larvae with *M. mycetomatis*, extracted total RNA at 0h (healthy larvae), 4h, 24h, 72h and 168h post-infection (**Figure 2A)** and performed RNA-Seq and LQ-ssCAGE and profiled their transcriptomic responses at each time point. Based on the grain size, the total number of grains and the melanisation of the hemolymph, the maximum burden of the infection was observed 72 hours after infection (**Figure 2B**). All larvae died within 192 hours after infection (**Supplementary Figure S2**). During infection, mycetoma grains developed over time. Early grains, which were small in size and loosely structured, were observed at 4 hours post-inoculation. After 24 hours, more mature grains were observed. The hyphae formed within the grain and the immune cells (hemocytes) were present in the cement material. At the maximum burden of infection, mature grains were present and surrounded by a capsule and immune cell (**Figure 2C**). In the late grain, present at 168 hours, disintegration of both the capsule and grain was observed, and immune cells were entering the grain.

We noted that 88.8% of the total RNA-Seq reads in the grains originate from the *G. mellonella* larvae (Host), while only 0.01% of the RNA-Seq reads originate from *M. mycetomatis* (Pathogen) (**Supplementary Table S1**). The rest of the RNA-Seq reads (11.19%) are unmapped reads. Likewise, for LQ-ssCAGE, we found that 31.6% of the reads are mapped to the *G. mellonella* larvae, while only 0.66% reads mapped to the *M. mycetomatis* (**Supplementary Table S1**). This was not unexpected. It has already been demonstrated that in a typical mycetoma grain, only roughly 10% of *M. mycetomatis* DNA was found, the remaining DNA originated from the host (20).

The t-*SNE* clustering of the host and pathogen transcripts mapped to the *G. mellonella* genome shows strong clustering of the samples per each time point (**Figure 2D**). The three samples of the cultured *M. mycetomatis* pathogen (yellow dots) cluster separately from the rest of the samples (host). RNA-Seq and LQ-ssCAGE expression levels of *G. mellonella* larvae genes positively correlated across all samples, with a median Spearman’s correlation (0.51) (**Figure 2D**). The t-*SNE* plot in (**Figure 2D**) indicates the quality of the transcriptomics profiling. The *G. mellonella* reads mapped to the pathogen genome indicate the presence of the conserved reads between the two species (host and pathogen) and the presence of pathogen reads after infection (**Figure 2D**).

### *G. mellonella* promoters and active enhancers expression landscape during infection

LQ-ssCAGE enabled mapping of the transcription start sites (TSS) and, therefore, the regulatory elements (Promoters and active enhancers) (21). The mapped TSS of the *G. mellonella* larvae libraries were used for identifying promoters and active enhancers (**Material and Methods**). Based on the CAGE tags generated by LQ-ssCAGE, we defined (16,548) expressed TSS from all libraries (n=16). We found that 43.1% (7,129) of the defined *G. mellonella* TSS are promoters followed by 15.1% (2,495) Coding Sequence (CDS), 13.3% (2,209) intergenic and 10.5% (1,731) intron (**Figure 3A**). Additionally, LQ-ssCAGE enabled the identification of 11,031 active enhancers in *G. mellonella*. Of these defined bidirectional transcribed enhancers, 34.5% (6,466) were overlapping intronic regions, and 24.4% (4,565) were overlapping intergenic regions (**Figure 3A**). The principal component analysis of the defined *G. mellonella* LQ-ssCAGE TSS (**Supplementary Figure S3A**) separated the healthy larvae from the infected larvae samples and the samples per time point (grain development stage). We found that all defined TSS were expressed in healthy and infected larvae (**Figure 3A**).

To understand the role of differential expression of the TSS and active enhancers, we used edegR (**Material and Methods**). We observed that of the total 16,548 *G. mellonella* TSS, only 3,279 TSS were significantly differentially expressed (**Supplementary Table S2)**. The most differentially expressed TSS are down-regulated between T4h-T168h. A summary of the up and down-regulated TSS and active enhancers is shown in (**Supplementary Figure S3B**).

To identify potential transcription factor binding sites (TFBSs) in the *G. mellonella* larvae genomes, we used HOMER Motif tools (**Material and Methods**) (22). Using the promoter regions, we predicted 28 transcription factors. We performed motif activity analysis to analyse trans-regulatory elements that regulate distant genes. The motif activities (**Figure 3B**) represent the average expression level of genes with a predicted binding site for each motif. **Figure 3B** shows the top 10, broadly expressed transcription factors (TF). The TFs in **Figure 3B** are ordered by the -log p-value, as an example, the TF LOC113521785 (iroquois-class homeodomain protein IRX-4-like) and (LOC113511740) GATA-binding factor C are highly expressed in at T4h (pre-mature grain stage). The gene ontology (GO) biological process associated with these genes is the regulation of transcription. The motif activity analysis indicates that at the time of the infection of the larvae, TFs altered their activity based on the eumycetoma grain development stage. A list of all predicted TF and associated GO terms is provided in **Supplementary Table S3**

### *G. mellonella* differentially expressed genes (DEGs) exhibited grain development stage-specific expression

From the time-course transcriptomic profile and analysis using the RNA-Seq data (**Material and Methods**), we observed that DEGs manifested in eumycetoma grain development stage-specific expression. We investigated two types of DEGs. First, the sharp changes, which detect the changes between pre-infection and post-infection. Second, the consecutive changes, which detect the changes between consecutive time-points, and between every time point and 0h (**Material and Methods**). To understand the patterns of the expression of the DEGs per each time-point (grain development stage), we performed hierarchical clustering of the top 50 DEGs. The clustering shows changes of expression of the top 50 DEGs (**Figure 4A**), at 0h vs. 4h, most of the genes become upregulated at 4h. In 0h vs. 24h, most of the genes become downregulated after 24h. Similar patterns were observed for 0h vs. 72h and 0h vs. 168h. Likewise, we observed the changes in DEGs in the rest of the time points 4h vs. all, 24h vs. all and 72h vs. 168h (**Supplementary Figure S4**). Validation by RT-qPCR of selected differentially expressed genes with known functions, in the infected *G. Mellonella* libraries confirmed their expression in these samples (**Supplementary Table S4** and **Supplementary Figure S5**).

**Figure 4:**
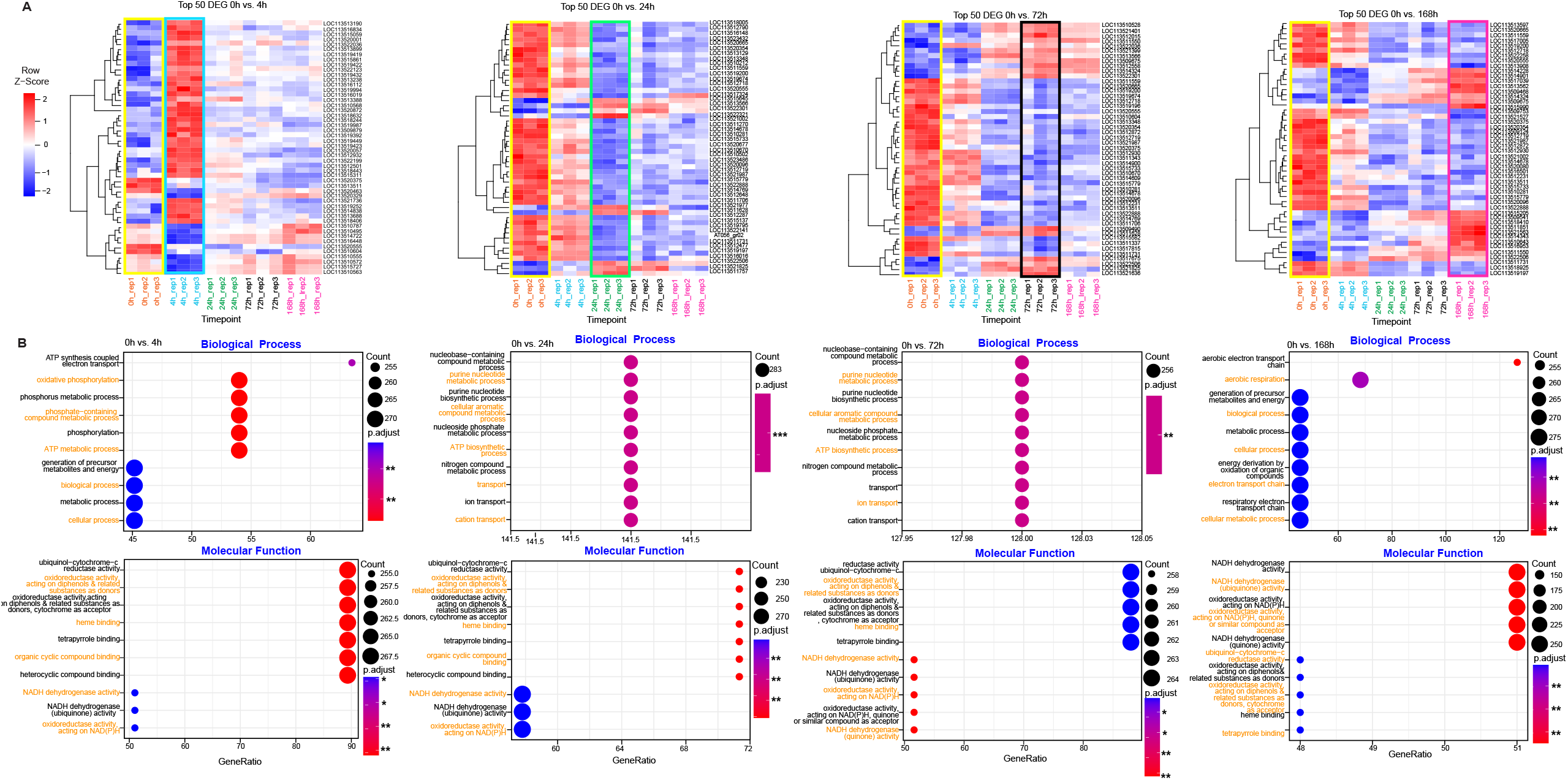
Hierarchical clustering of the host top DEGs and gene set enrichment analysis. **A**. Heatmaps showing the hierarchical clustering of the top 50 DEGs of *G. mellonella* for four consecutive changes (**Material and Methods**). Columns represent the replicates, and rows represent the host gene symbols. Gene names are provided (**Supplementary Table S5**). Replicates are colored and grouped per time point. 0h replicates are highlighted with yellow rectangles. Heatmap cells colored by Z-score scale (−2 to 2). **B**. Dotplots showing enriched pathways in biological process and molecular functions in the host during infection. Each pair of plots shows enriched pathways of the top DEGs in each consecutive point. Each pathway is represented as a dot. The dots are colored by the p-adjusted value as computed by gseGO function of clusterProfiler R package *P < 0.05, **P < 0.01, ***P < 0.001, ****P < 0.0001. The count represents the number of genes that belong to a given gene-set, and the GeneRatio represents the count/setSize. setSize is the total number of genes in the gene-set. Each pathway corresponds to a Gene Ontology (GO) term.

The Venn Diagram in (**Supplementary Figure S6 A**) summarized the consecutive changes in DEG. For the In sharp change category, 4,338 genes are differentially expressed between the healthy and infected larvae (**Supplementary Figure S6 B**). We summarized the DEGs in all consecutive changes and the sharp changes. We found that 3,575 genes are differentially expressed (**Supplementary Figure S6 C**). The Volcano plot in **Supplementary Figure S7** summarized the p-values and fold change of the DEG per time point.

### Gene set enrichment analysis of the top *G. mellonella* DEG

To understand the biological meaning of the DEG, we performed a gene set enrichment analysis of the top DEG (**Material and Methods**). The enrichment analysis shows that several DEGs are significantly enriched in biological processes and molecular functions (**Figure 4B**). Several GO terms are enriched at each time point. At 0h vs 4h, most of the enriched GO terms are linked to oxidative phosphorylation (GO:0006119), ATP metabolic process (GO:0046034), energy derivation by oxidation of organic compounds (GO:0015980), generation of precursor metabolites and energy (GO:0006091). All these terms are linked to energy pathways. The 0h vs 24 and 0h vs. 72 were mostly enriched in the nucleobase metabolic process, purine nucleotide biosynthetic process, and ion and cation transport. Among the latter, genes linked to iron homeostasis such as melanotransferrin, cytochrome b561-like isoforms, aconitate hydratase and proton-coupled folate transporters were differentially expressed. At the late-stage grain development (0h vs 168h), mostly enriched GO terms are related to aerobic electron transport chain and energy pathways. The full list of the top 50 DEGs and the associated annotation is in **Supplementary Table S5**.

### *M. mycetomatis* genes are differentially expressed in infected *G. mellonella* larvae

To study the presence and expression of the pathogen genes in infected *G. mellonella* larvae, we mapped all samples to the *M. mycetomatis* genome assembly ASM127576v2 (**Material and Methods**). We observed the same time-patterns of expression as in the expression of the host DEGs, when we clustered the top 20 pathogen DEGs (**Figure 5A**). Genes which appeared to be differentially expressed at all time points include Glyceraldehyde-3-phosphate dehydrogenase, 30kDa heat shock protein and ECM33, for which the corresponding proteins were previously demonstrated to be differentially abundant by LC-MS/MS (15). Complete annotation of the pathogen top 20 DEGs is provided in **Supplementary Table S6**. The GSEA of the top DEG of the pathogen is shown in **Figure 5B**, with several GO terms enriched.

**Figure 5:**
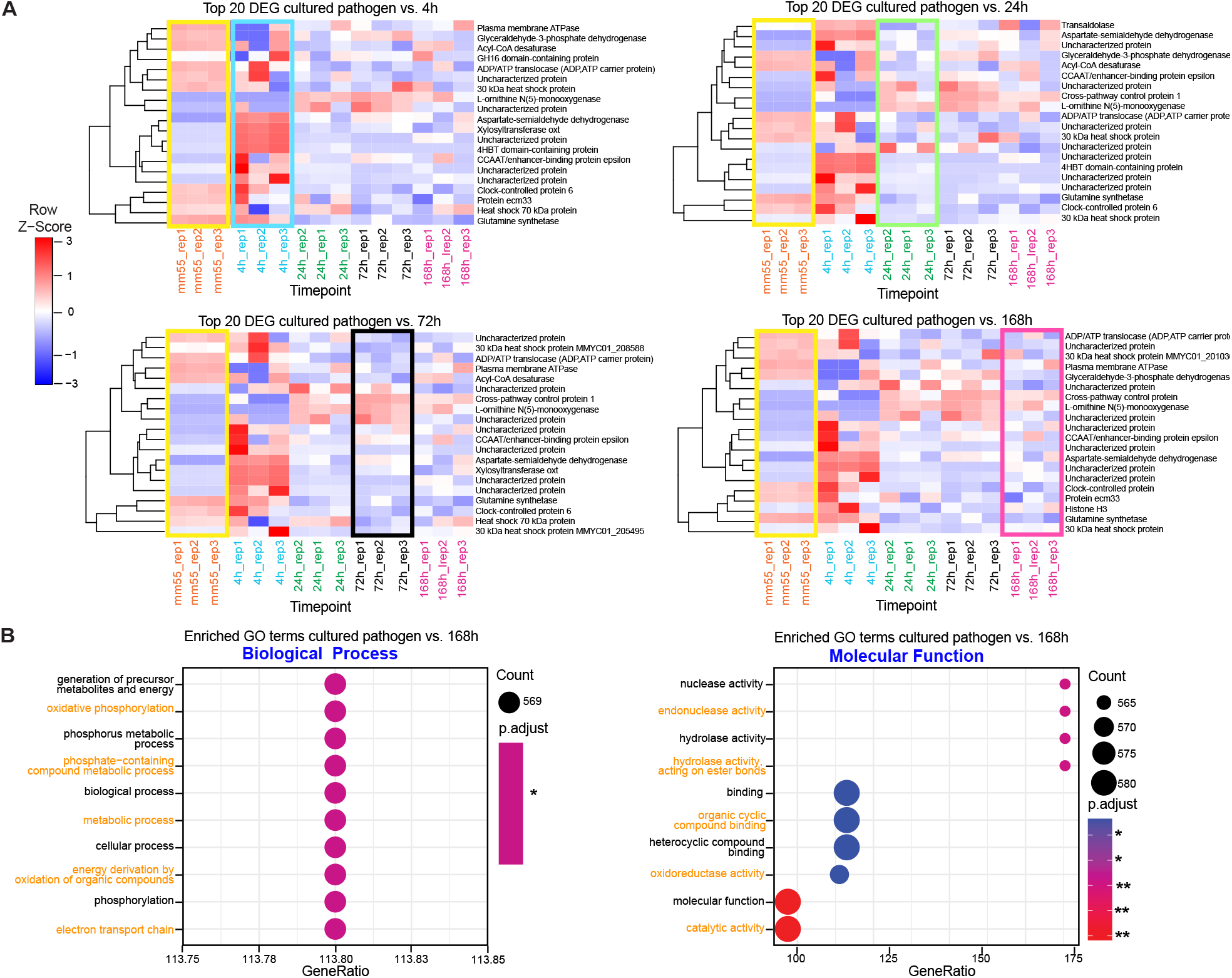
Pathogen reads in infected host and gene set enrichment analysis. **A** Heatmaps of Top 20 DEG cultured pathogen vs. pathogen reads in infected host 4h. Full annotation of the genes in each of the heatmaps in (**Supplementary Table S3**). **B**. GSEA of the top 20 DEG pathogen genes and enriched GO terms. The dots are colored by the p-adjusted value as computed by gseGO function of clusterProfiler R package. *P < 0.05, **P < 0.01, ***P < 0.001, ****P < 0.0001.

### Host and pathogen transcript dynamic and response patterns

Since we ended up with high number of DEGs from the RNA-Seq data, we further investigated the dynamic of the expression and response pattern of the host and pathogen transcriptome to the infection separately (**Figure 6A**). For the host DEGs (3,498), we observed a set of host genes start to gradually respond to the infection by either upregulation or downregulation after 0h. We classified those genes as gradual response genes. The second set of DEGs were classified as early response genes, which immediately change the expression pattern upon infection. Most of the genes in this category change their expression either up or down at 24h. The third set of DEGs were the early-long response patterns in which the immediate change in transcription at 4h-24h remains longer during the course of infection. The last pattern of the response was the late response genes, these genes start to change their expression at 24h. **Figure 6B** shows an example of a gene for each expression pattern category. Detailed annotation of genes in each response pattern was obtained from the UniProt database (**Supplementary Table S7**) (23). For the pathogen genes, we observed all the above patterns as in the host, except the late response pattern genes. Of the total 136 pathogen DEGs, only nine genes gradually change expression after the infection (**Figure 6C**). Interestingly, among these genes, Ecm33 and the stress-related hsp30 were noted (24). Proteomic data demonstrated that protein Ecm33 was present in the mycetoma grain at all time points analyzed (15). Bar plots in **Figure 6D** demonstrate the expression of example genes. Detailed annotation of the 136 pathogen genes with corresponding GO term annotation is listed in **Supplementary Table S8**.

**Figure 6:**
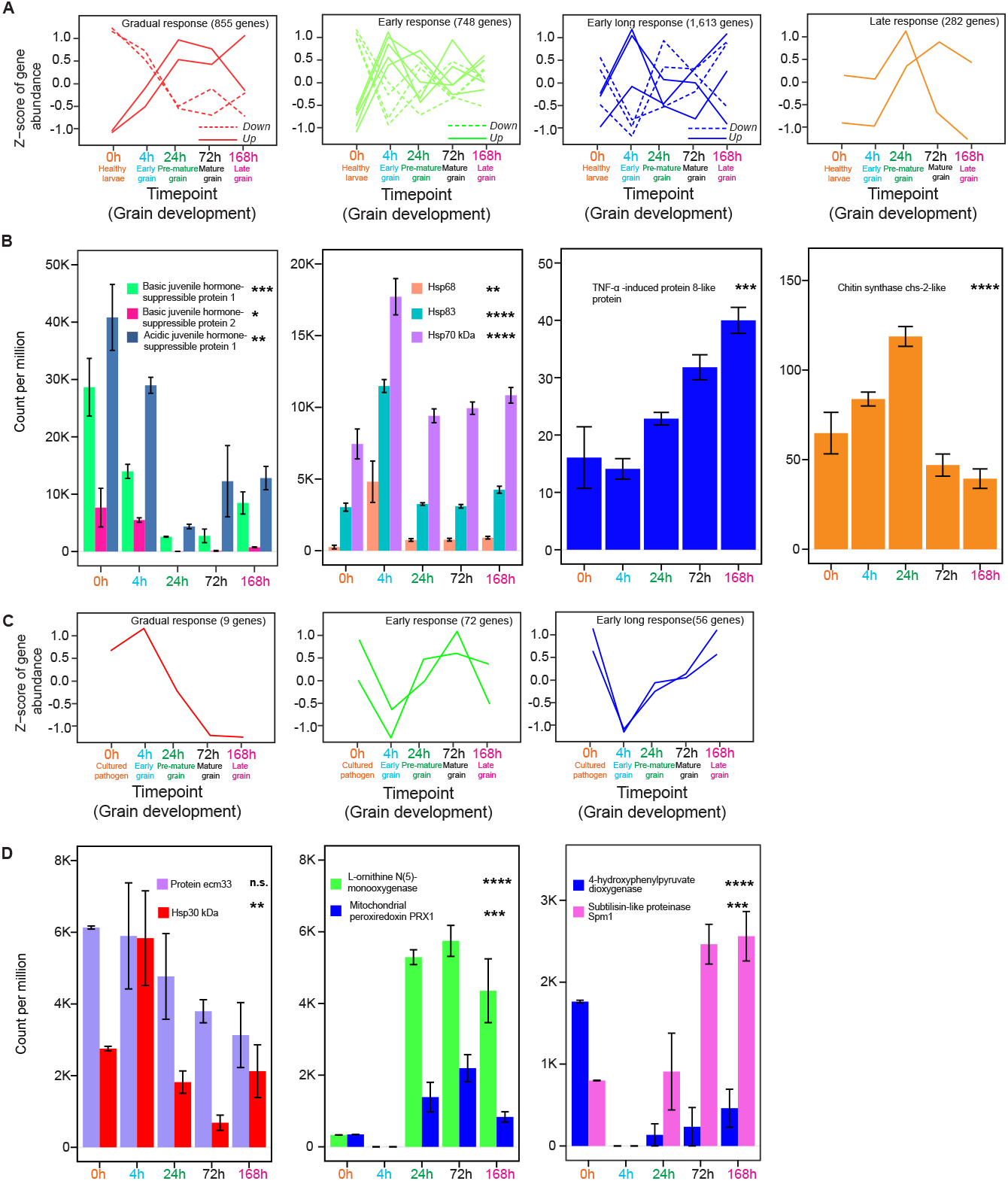
Host and pathogen transcriptomic changes before and after the development of Eumycetoma grain. **A**. The responses of host DEGs are shown in four patterns (Gradual, early, early long, and late response). Y-axis represents the Z-score of gene abundance, and the X-axis shows the time point and grain development stage. The gradual, early response and early long response show sub-patterns (down and up) shown as dashed or continuous lines respectively. Full annotation of the genes in each response group in (**Supplementary Table 6**). **B**. Expression of some of the host DEGs corresponding to the patterns in (A). Barplot with standard error-of-mean (SEM), Y-axis shows the count per million (CPM), and X-axis shows the time point. **C**. Transcriptomic response patterns of the pathogen DEGs. Only three patterns are detected. As in A, Y-and-X axis represent the Z-Score gene abundance and grain development stage, respectively. Full annotation of the genes in each response group in **Supplementary Table S8. D**. Expression of some of the pathogen DEGs corresponding to the patterns in C. **\***P < 0.05, **P < 0.01, ***P < 0.001, ****P < 0.0001, and n.s., not significant by two-way ANOVA test; Data in the bar chart are represented as expression ± SEM.

### Genes involved in iron sequestering are differentially expressed and contribute to eumycetoma grain development

To investigate the biological relevance of the differentially expressed genes, we performed Protein-Protein Interaction (PPI) network enrichment analysis of relevant genes. We first decided to focus on biological processes known to be expressed by *M. mycetomatis*. We demonstrated that melanisation of the grains occurs rapidly during the course of infection, and some of the genes involved in the melanin biosynthesis pathway were differentially expressed (e.g., 4-hydroxyphenylpyruvate, **Figure 6D**). Based on this information, we expected the complete biosynthesis pathway for DHN-melanin and pyomelanin to be expressed during infection. For DHN-melanin and pyomelanin, NCBI blast found significant similar homologous in *M. mycetomatis* for all genes. We identified all homologues of *A. fumigatus* DHN-melanin and pyomelanin biosynthesis pathways in *M. mycetomatis* using the NCBI Blast web service (**Supplementary Table S9**). **Supplementary Figure S8** shows the DHN-melanin biosynthesis pathway and pyomelanin biosynthesis pathway in *A. fumigatus* and the PPI networks of homologous genes in *M. mycetomatis*. These PPIs indicate the functional interaction link between genes involved in both the DHN-melanin and pyomelanin biosynthesis pathway and other genes in *M. mycetomatis* with a PPI network enrichment *p-value* of 3.08E-13 and 0.00388, respectively. The list of homologues of *A. fumigatus* DHN-melanin and pyomelanin in *M. mycetomatis* with their statistical report from NCBI blast is listed in **Supplementary Table S9**.

One of the main findings of the DGE analysis was significant expression changes in genes related to heme-binding and iron regulation. Therefore, we decided to further analyse these regulatory pathways for both *G. mellonella* and *M. mycetomatis*.

To investigate the iron regulation response of *G. mellonella* during infection, we focused on genes encoding for expression of ferritin and transferrin, proteins responsible for the transport and storage of iron (25). The list of homologues involved in iron regulation in *D. melanogaster* and *G. mellonella* with their statistical report from NCBI blast is listed in **Supplementary Table S10**. We constructed a PPI network, indicating the functional interaction of iron regulatory genes in *D. melanogaster* (**Figure 7A**). With a PPI network enrichment *p-value* of 0.00312, strong functional interaction is indicated between the following genes: Ferritin 1 Heavy Chain Homologue (Fer1HCH), Ferritin 2 Light Chain Homologue (Fer2LCH), Transferrin (Tsf1), Iron regulatory protein 1A (Irp-1A), Iron regulatory protein 1B (Irp-1B) and Malvolio (Mvl). Except for Mvl, homologues of the respective genes were identified in *G. mellonella*, and expression levels are depicted in the bar chart (**Figure 7A**). The genes ferritin subunit (LOC113510018), ferritin lower subunit (LOC113510017), transferrin (LOC113509694), aconitate hydratase (LOC113516537/ LOC113511518) and cytoplasmic aconitate hydratase (LOC113522652), are all consistently expressed in both healthy and infected *G. mellonella* larvae over time. This data is in line with previous literature in which mRNA expression of ferritin in healthy *G. mellonella* larvae is detected, and mRNA expression of transferrin is detected in both healthy larvae and infected larvae of *Mandunca sexta* (26, 27). In contrast with a proteomic analysis of healthy G. mellonella larvae and larvae infected with *M. mycetomatis*, we did not observe differential expression of both ferritin and transferrin coding genes (15).

**Figure 7:**
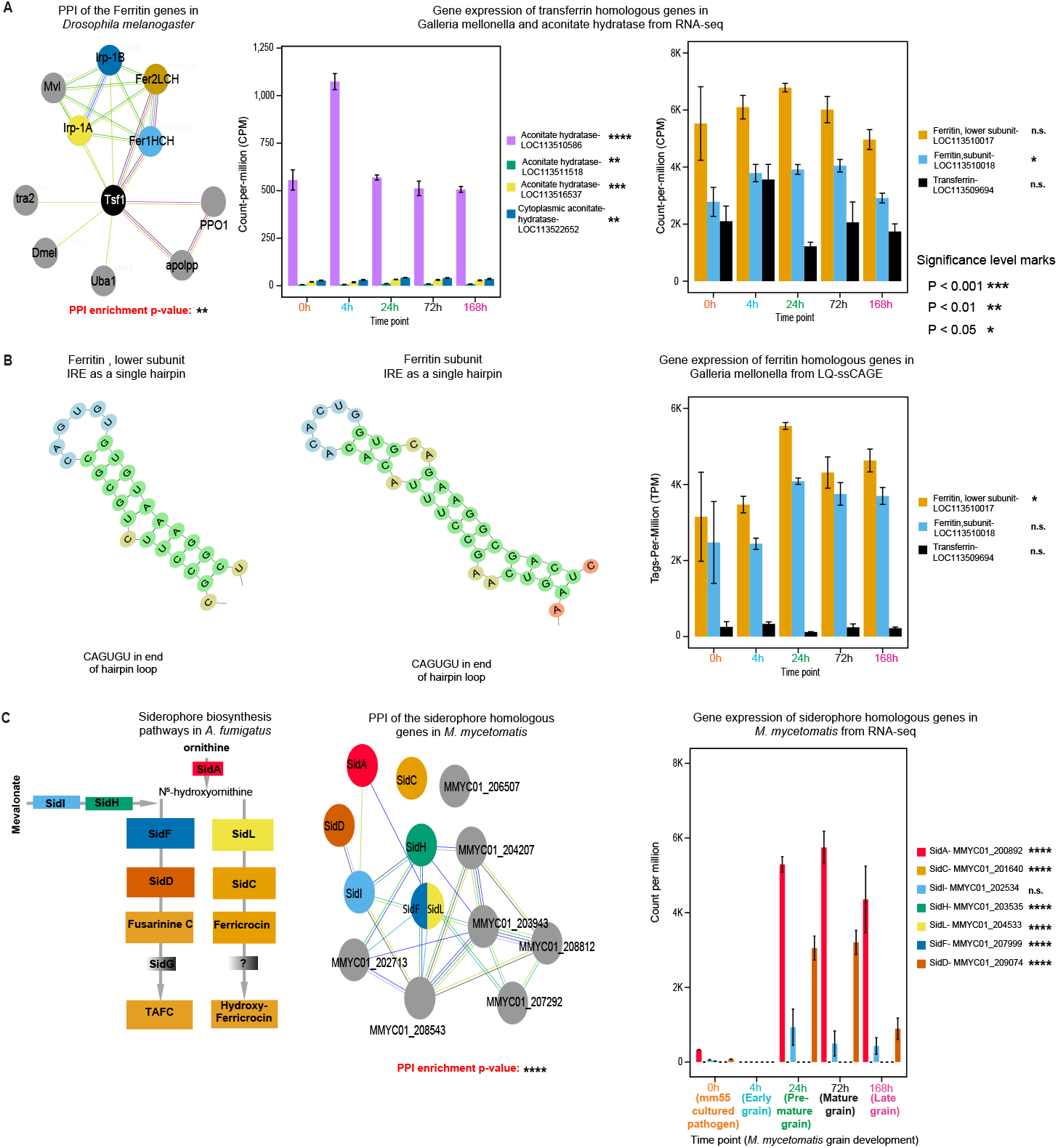
Iron regulation in *G. mellonella* and siderophore biosynthesis pathways in *M. mycetomatis*. **A**. PPI of Ferritin genes in *Drosophila melanogaster* adapted from G. Xiao et al., with a focus on transferrin (Tsf1), Ferritin 1 Heavy Chain Homologue (Fer1HCH), Ferritin 2 Light Chain Homologue (Fer2LCH), Iron regulatory protein 1A and 1B (Irp-1A and Irp-1B) (49). Expression levels of respective *G. mellonella* homologues during infection are shown in the bar plot. Aconitate hydratase encoding genes LOC113511518, LOC113516537, LOC113522652 and LOC113510586, are differentially expressed during infection. **B**. Generated hairpin loops in LOC113510017 and LOC113510018 transcripts, containing the conserved CAGUGU sequence characteristic for Iron Responsive Elements (IREs). Promotor and active enhancer activity of ferritin and transferrin homologous genes in *G. mellonella* based on LQ-ssCAGE. **C**. Siderophore biosynthesis pathway adapted from Gründlinger et al. (32). The pathway starts with mevalonate converted to anhydromevalonyl-CoA by SidI and SidH (33). A. fumigatus uses three hydroxamate-type siderophores for iron uptake. The extracellular triacetylfusarinine C (TAFC), hyphal ferricrocin (FC) and conidial hydroxyferricrocin (HFC) (33). N□-hydroxyornithine generated from ornithine by SidA. The SidG gene has no homologues gene in *M. mycetomatis*, and the gene marked as (?) is not characterized yet in *A. fumigatus*. In the PPI, SidF and SidL are annotated to the same gene in *M. Mycetomatis* (N(6)-hydroxylysine O-acetyltransferase). No significant similarity was found for SidG in *M. mycetomatis*. All siderophore genes except SidG are differentially expressed in RNA-Seq data, as shown in the bar plot. *P < 0.05, **P < 0.01, ***P < 0.001, ****P < 0.0001, and n.s., not significant by two-way ANOVA test; Data in the bar chart are represented as expression ± SEM.

Gene expression does not always directly correlate to protein expression. Therefore, we postulated that the presence of one or multiple translational regulatory mechanisms might explain the difference between the proteomic and transcriptomic findings. This is strengthened by our PPI analysis and gene expression data (**Figure 7A**), showing the expression of multiple aconitate hydratases (Irp-1A and Irp-1B), genes encoding for iron-regulatory proteins which regulate translation by binding to iron-responsive elements (IRE) in the 5’ UTR region (28). Of the multiple aconitate hydratases, homologues expressed, LOC113511518, LOC113516537, LOC113522652 and LOC113510586 were differentially expressed (**Figure 7A**). To further validate the functional similarity with known iron regulatory processes, we identified the IRE regions present in mRNA transcripts. These IRE regions manifest in the form of a hairpin loop, containing a conserved CAGUGU at the end of the hairpin loop. **Figure 7B** shows the identified IRE hairpin structures of the ferritin subunit and ferritin lower subunit in the 5’ UTR region. The IRE of the lower ferritin subunit was identical to the one earlier described for the 32 kDa *G. mellonella* subunit, while the IREs for the upper ferritin subunit differed in a few amino-acids from the one described for the 26 kDa ferritin subunit (29, 30). LQ-ssCAGE data reveals expression activity of the promoter regions for the ferritin subunit (LOC113510018), ferritin lower subunit (LOC113510017) and transferrin (LOC113509694) (**Figure 7B**), indicating that these regions were actively expressed. Altogether, this data supplements the proteomic findings, suggesting the presence of an iron regulatory mechanism in *G. mellonella* for the overexpression of ferritin to store intracellular iron molecules during *M. mycetomatis* infection.

In response to the withdrawal of iron from haemolymph by *G. mellonella, M. mycetomatis* needs to alter its iron regulation. Among the top 20 DEG for *M. mycetomatis*, L-ornithine N(5)-monooxygenase (homologues to SidA) and non-ribosomal peptide synthetase SidD were identified (**Figure 5A**), which are key in the siderophore biosynthesis pathway found in *A. fumigatus* and *A. nidulans* (**Figure 7C**) (31, 32). NCBI blast found significant similar homologues in *M. mycetomatis* for all genes except SidG (**Supplementary Table S9**). PPI analysis of relevant genes for the *M. mycetomatis* homologues was performed. The PPI network analysis (**Figure 7C**) revealed a strong interaction between the expressed *M. mycetomatis* homologues for the entire siderophore biosynthesis as described for *A. fumigatus* (32). The PPI network obtained with enrichment *p-value* (1.83E-14) showed no interaction between SidC and the rest of the network. In the PPI network (**Figure 7C**). The top enriched biological processes (Gene Ontology) are GO:0006635 (Fatty acid beta-oxidation), GO:0006631 (Fatty acid metabolic process), GO:0003995 (acyl-CoA dehydrogenase activity), GO:0003857 (3-hydroxyacyl-CoA dehydrogenase activity). The top KEGG pathways enriched in the PPI network are map01100 (Metabolic pathways) FDR (1.11E-05) and map00071 (Fatty acid degradation) FDR (0.0024). Both Acyl-CoA ligase sidI and Mevalonyl-coenzyme A hydratase sidH are the third-degree hub nodes in the PPI network (**Figure 7C**). In *A. fumigatus* SidI and SidH linking biosynthesis of mevalonate and triacetylfusarinine C (TAFC) (33). Seven genes found to have functional link in the PPI network in (**Figure 7C**) (grey nodes), these genes are MMYC01_206507 (5’-hydroxyaverantin dehydrogenase), MMYC01_204207 (3-hydroxyisobutyryl-CoA hydrolase), MMYC01_203943 (Short/branched chain specific acyl-CoA dehydrogenase), MMYC01_208812 (Glutaryl-CoA dehydrogenase), MMYC01_202713 (3-hydroxybutyryl-CoA dehydrogenase), MMYC01_207292 (Acetylglutamate kinase), and MMYC01_208543 (3-hydroxybutyryl-CoA dehydrogenase). All siderophore homologous genes in *M. mycetomatis* are differentially expressed. We observed that L-ornithine N(5)-monooxygenase (SidA), Acyl-CoA ligase (SidI) and Nonribosomal peptide synthase (SidD) are highly expressed in all timepoints with significant upregulation one day after the infection compared to the rest of the genes involved in the siderophore pathway (**Figure 7C)**.

## Discussion

In this transcriptomic study, we demonstrated that 3,498 *G. mellonella* and 136 *M. mycetomatis* genes are differentially expressed during the course of grain formation, and the extensive data set generated during this study can be used to interrogate various processes important in grain formation.

In this paper, we focused on the process of iron homeostasis. In both host and pathogen, we found numerous genes involved in this process to be differently expressed.

To limit the access of iron for pathogens such as *A. fumigatus, Cryptococcus neoformans* and *Mycobacterium tuberculosis* (34, 35, 36) is an important innate defense mechanism from the host. This process, also known as nutritional immunity (37), is most studied in mammals, little in insects, and hardly in *G. mellonella*. Therefore, our data adds to the knowledge of the iron response in *G. mellonella* during infection. Generally, in insects, iron sequestering, transport and storage are regulated by both ferritin and transferrin (25, 38, 39).

In our study, we did find expression of the ferritin subunit (LOC113510018), ferritin lower subunit (LOC113510017) and transferrin (LOC113509694) at all time points investigated. However, in contrast to what was seen in other fungal infections, expression levels did not significantly alter during infection (40). Transferrin and ferritin protein levels previously measured in the same *G. mellonella* infection model, on the other hand, were differentially abundant. In that study, a 5 to 12-fold decrease in transferrin protein levels and a 3-fold increase in ferritin protein levels were noted during *M. mycetomatis* grain formation in *G. mellonella* (15). For ferritin, this difference can be explained by the process of iron regulation via iron regulatory proteins (IRP). In *G. mellonella*, these IRPs are encoded by aconitate hydratases, and *G. mellonella* aconitate hydratases LOC113516537, LOC113511518 and LOC113522652 were differentially expressed during *M. mycetomatis* infection. IRPs bind IREs under low iron conditions to repress the translation of ferritin. A process well studied in mammals (30, 41). In mammals, IRPs respond to low iron by binding to IREs present in the 5’UTR of the mRNAs of ferritin (37). This results in translational repression of ferritin. In contrast, in the presence of enough iron, the enzyme aconitate hydratase will form a [4Fe-4S] cluster, which prevents the IRP binding to the IRE. This in turn, results in translational activation of ferritin (41). Since we and others identified IREs in the 5’UTR regions of ferritin from *G. mellonella* and in *D. melanogaster*, and it was described that IRP1 was able to bind to the ferritin IRE to regulate translation, we postulate that in *G. mellonella* the regulation of ferritin is similar (**Figure 8**) (28, 29, 30).

**Figure 8:**
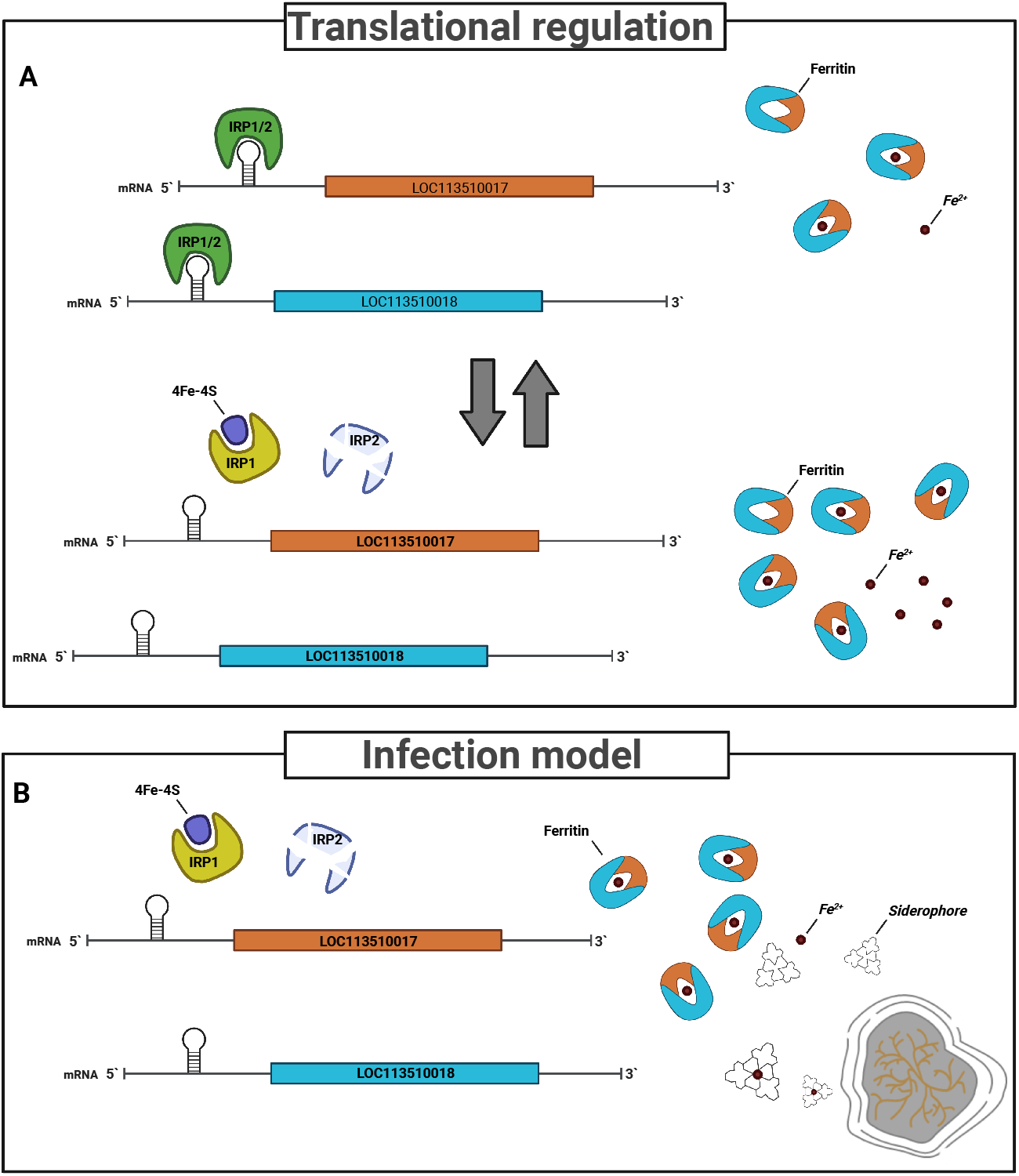
Proposed model of Iron regulation in *Galleria mellonella* during infection with *M. mycetomatis*. **A**. In healthy larvae, in the absence of iron (Fe^2+^), the IRP1 and IRP2 form a complex, which can bind to Iron Responsive Elements (IRE) in the 5’ prime UTR region of the ferritin subunits coding transcripts LOC13510017 and LOC13510018, preventing translation of the mRNA to form ferritin. In the presence of iron (Fe^2+^), 4Fe-4S is formed, binding to IRP1. The IRP1/4Fe-4S complex prevents binding to the IRE, enabling translation of both LOC13510017 and LOC13510018, forming ferritin. Ferritin will bind Fe^2+^ for transportation and storage. **B**. During infection, increased expression of aconitate hydratase results in increased formation of the 4Fe-4S complex, binding of IRP1 and thus the translation of both subunits of ferritin for the binding of Fe^2+^, creating an iron-derived environment. In response to the limited available iron, the fungus shows significantly increased expression of the siderophore-related biosynthesis pathway for the production of siderophores needed for sequestering Fe^2+^, which is essential for the survival of the fungus.

Due to the lack of IRE in the transferrin UTR regions, the discrepancy between transferrin expression levels and protein levels cannot be due to IRP/IRE regulation. In other pathogenic fungi, such as *Candida albicans* and *A. fumigatus*, it has been demonstrated that mammalian transferrin can be used as an iron source for the fungus (42, 43). *C. albicans* forms a complex with transferrin and obtains iron via the high-affinity reductive system (42). In *A. fumigatus*, siderophores play a role in the removal of iron from transferrin (43). The time needed for *A. fumigatus* to produce enough siderophores to remove iron from transferrin was eight hours, however, 22 hours were needed to degrade the complete transferrin protein (43). In our larval model, it took 24 hours to get a 5-fold decrease in transferrin protein abundancy and 72 hours to get a 12-fold decrease in transferrin protein abundancy in the mycetoma grain (15). Furthermore, in the current study, we also demonstrated that in our *M. mycetomatis* larval infection model, the genes involved in the siderophore biosynthetic pathway were differentially expressed during infection. After 24h infection, a strong increase in the expression of especially SidA, SidD and SidI was observed (**Figure 7**), coinciding with the decrease in transferrin protein abundancy; this indicates that a similar system could exist in *M. mycetomatis* during infection.

Next to obtaining iron from transferrin, fungal siderophores can also obtain iron from other sources surrounding the fungus. Our PPI analysis demonstrated that almost all homologues of the *A. fumigatus* siderophore biosynthetic pathway were present in *M. mycetomatis*. Only the homologue of the *A. fumigatus* gene encoding for FsC-acetyl coenzyme A-N(2)-transacetylase (SidG), the final enzyme required for the production of the *A. fumigatus* siderophore triacetylfusarinine C (TAFC), was not found in the *M. mycetomatis* genome. SidG is only required for the biosynthesis of TAFC, not for the siderophore fusarinine C (**Figure 7C**), therefore, it is not surprising that there was no homologue in *M. mycetomatis* (44). So far, homologues of SidG have only been found in a few *Aspergillus, Fusarium* and *Nectria* species, but not in species lacking TAFC. Based on this, it is most likely that *M. mycetomatis* produces an extracellular fusarinine C-type siderophore and one or two intracellular ferrichrome-type siderophores, such as ferricrocin and hydroxyferricrocin. The presence of hydroxamate-type siderophores, which are often composed of trans-fusarinine units, has been demonstrated to be produced by *M. mycetomatis* previously, however they were never chemically 14haracterized (45).

The exact role of the acquisition of iron via transferrin or siderophores in the pathogenicity of *M. mycetomatis* has not been described. However, in *A. fumigatus*, it has been demonstrated that the elimination of the entire siderophore biosynthesis resulted in avirulence in a murine model of invasive aspergillosis (46, 47). Furthermore, it has also been described that siderophores can be used either diagnostically or therapeutically (48). Therefore, additional studies on the role of siderophore biosynthesis in *M. mycetomatis* could improve the management of mycetoma in the future.

In this study, we used a transcriptomic approach to decipher the response of the *G. mellonella* host and the *M. mycetomatis* pathogen in mycetoma grain formation. Although a large number of genes were differentially expressed, we demonstrated that iron metabolism plays an important role in *M. mycetomatis* grain formation in *G. mellonella* over time during infection. The data generated in this study is of enormous value to the scientific community and can be re-used to answer other biological questions. The identification of the importance of iron acquisition during grain formation can be exploited as a potential novel diagnostic and therapeutic strategy for mycetoma.

## Contributors

Conceptualization: IA, WVD

Methodology: IA, MK, WL, WVD

Library preparation: RIM, TK

Raw sequence processing IA, AH, CT, MT

Investigation: IA, MK, WVD

Visualization: IA, MK, WVD

Cultured pathogen: AHF

Supervision: YO, AV, TK, IA, WVD

Writing—original draft: IA, MK, WVD

Writing—review & editing: IA, MK, AV, AHF, TK, WVD

## Acknowledgments

We greatly appreciate the efforts of Atsui Hiroto, Nobuyu Takeda, Teruaki Kitakura, and Akira Furukawa in providing technical support. We are thankful for the English proofreading by Scott Walker, and Hiroko Kinoshita for her helpful support for LQ-ssCAGE library preparations. We thank RIKEN Center for Integrative Medical Sciences sequence platform for their helpful support to sequence RNA-Seq and LQ-ssCAGE libraries.

This work was supported by research grants from Dutch Research Council by Aspasia grant no. 015.013.033 and by the Erasmus University with an EUR Fellowship, both awarded to Wendy W.J. van de Sande. This study is also supported by Research Grants from the Japanese Ministry of Education, Culture, Sports, Science and Technology MEXT to RIKEN Center for Integrative Medical Sciences.

## Data Sharing Statement

Raw sequence data and expression tables usedtables used in the analysis of this article have been 570depositeddeposited in the Gene Expression Omnibus (GEO) repository, https://www.ncbi.nlm.nih.gov/geo571, https://www.ncbi.nlm.nih.gov/geo (accession numbers: <u>GSE213321</u>, <u>GSE213322</u>, <u>GSE213329</u>, and <u>GSE213332</u>).

