## Supporting_Information_Methods_Figures for "Iron regulatory pathways differentially expressed during *Madurella mycetomatis* grain development in *Galleria mellonella*"

### **Supporting Information Text**

#### Analysis of the fungal burden in *G. mellonella*

The fungal burden was determined by manually counting the grains using 40x magnification of a light microscope mounted with a Canon EOS70D camera (Canon Inc). Counting was performed by three independent scientists. The grains were visualized on a computer screen using EOS Utility software (Canon Inc.) and categorized into large (>0.02mm^2^), medium (0.01-0.019 mm^2^) and small (0.005-0.009 mm^2^) sizes as described by Lim and colleagues (1). The sum of all grains represents the total amount of grains within the larvae. The total grain size within the larvae was determined by multiplying the sum of all grains with the minimum size of their respected categories. The difference in the total number of grains or the total grain size observed on the respective time points were determined using the Mann-Whitney *U* test in GraphPad Prism 8. A p-value > 0.05 was deemed significant.

#### Library preparation, sequencing, mapping, and processing

##### ***RNA-seq***

HiSeq2500 instrument from Illumina was used for sequencing with Paired-End; 100base followed by base calling. The quality of the raw reads was assessed using FastQC [https://www.bioinformatics.babraham.ac.uk/projects/fastqc/]. We returned 245,960,792 total tags for the *G. mellonella* larvae samples and 152,647,712 total tags for the *M. mycetomatis* samples.

*M. mycetomatis* raw reads mapped against *M. mycetomatis* genome assembly ASM127576v2 (2)*.* The *G. mellonella* larvae reads were mapped twice, first against *G. mellonella* assembly ASM364042v2 and then against *M. mycetomatis* genome assembly ASM127576v2*.* We used STAR Aligner version 2.7.1a with default settings (3). In short, for each of the *M. mycetomatis* libraries, on average, 43.8 million reads were mapped, of these, on average, 75.8% were uniquely mapped. For each of the *G. mellonella* larvae libraries, on average, 49.1 million reads were mapped. In the *G. mellonella* larvae libraries, 68,398 reads mapped to *M. mycetomatis* genome ASM127576v2.

##### ***LQ-ssCAGE***

LQ-ssCAGE sequencing libraries were prepared following the procedures described in (4). Like RNA-Seq, LQ-ssCAGE was prepared for both the *G. mellonella* larvae samples and *M. mycetomatis* with 25 ng as minimum total RNA input. The same HiSeq2500 instrument was used as described before but with Paired-End; 50base. After performing base calling for all samples, we returned 251,856,022 total tags for the *G. mellonella* larvae samples and 68,428,840 total tags for the *M. mycetomatis* samples. The same mapping strategy was used for LQ-ssCAGE as for RNA-seq described above. For each of the *M. mycetomatis* libraries, on average, 22.8 million reads were mapped, of these, on average, 35.0% uniquely mapped to the ASM127576v2 genome. For each of the *G. mellonella* larvae libraries, on average, 14.5 million reads were mapped, of these, on average, 47.0% were uniquely mapped against ASM364042v2.

##### ***Promoters and enhancers call***

To identify the promoters and enhancers for the two species, we created two Biostrings-based genome data packages. The two Bioconductor packages used by CAGEfightR to identify TSS and enhancers (5). The two packages were used for annotation of the CAGE-defined TSSs and gene-level expression. The mapped reads for each library were converted to CAGE tags (CTSSs) in BED format using an in-house script. The CTSS tag clusters (TC) were identified from the CTSSs and used for the subsequent analysis and quantification of the LQ-ssCAGE data. In short, for CTSS, counts were normalized using Tags-per-Million (TPM), and the normalized CTSS counts were used to calculate the pooled CTSS (CTSS signal across all samples). The pooled CTSS were further processed to reduce single tags spread across the genome. This is accomplished by removing CTSSs detected in only a single or few samples. The final set of the pooled CTSSs was used for the analysis at the level of clusters of CTSSs (Promoters and enhancers identification) and analysis at the level of annotated genes.

##### ***RNA-seq reads quantification***

To quantify the mapped reads from RNA-Seq libraries, the Rsubread FeatureCount function was utilized using the same parameters for RNA-Seq quantification (6). We tested the correlation between the RNA-Seq and LQ-ssCAGE (**Supplementary Figure S9**).

#### Exploratory analysis of RNA-Seq and LQ-ssCAGE data

The previously generated raw count matrix for RNA-Seq and LQ-ssCAGE was used for exploratory data analysis to explore the clustering of the samples per time point and expression patterns using R implementation of the t-distributed stochastic neighbour embedding (*t*-SNE) *t*-SNE. The Rtsne function of Rtsne package (V 0.15) with is_distance =TRUE and perplexity = 5. To explore the clustering of the host samples from the LQ-ssCAGE dataset mapped to the host genome, the prcomp function from stats (V 3.6.2) was used with standard settings and perform principle component analysis (PCA).

#### Prediction of the transcription factors binding sites (TFBS) and motif activity overtime

To predict transcription factors binding sites (TFBS) for the *G. Mellonella*, we used findMotifsGenome.pl from Homer. The input for the command is the set of identified Promoters and Enhancers from LQ-ssCAGE data. Also, the Homer command requires a FAST and GFF file for the *G. mellonella* genome, which was obtained from NCBI.

#### Differential expression analysis

To identify common DGE between consecutive and sharp changes, the result from the two steps regression analysis and the linear model analysis were further integrated. In short, a final LIMMA-derived *P* value was computed for each gene by combining all contrasts (e.g. T0-T4h) using eBayes function of LIMMA R package and computing the F-statistic, the associated *P* value, and the adj. *P* value using Benjamini-Hochberg multiple testing correction. A gene was differentially expressed if any of the following conditions applied: (a) LIMMA FDR *<* 0.001, (b) maSigPro R^2^ *>* 80%, or (c) LIMMA FDR *<* 0.01 & maSigPro R^2^ *>* 60%.

Results from the DGE analysis were visualized using a standard heatmap with customized settings for the heatmap.2 function from R package gplots (V 3.1.1). In addition to the heatmap, the EnhancedVolcano R package was used to plot the Volcano plots and visualize the results from the DGE.

#### Detect pattern of expression from DGE

To understand the expression pattern over time, the DEG from LIMMA and maSigPro of the RNA-Seq and LQ-ssCAGE data was analysed. This kind of analysis shows the gene expression dynamic per time point and the host response to the infection. The function degPatterns from the DEGreport R package (V 1.26.0) was used to cluster genes with similar expression pattern overtime. The input for degPatterns function expression matrix in logarithmic scale (only genes that are significantly different), the time course design experiment used to group samples (Healthy vs. infected larvae). In the degPatterns, the following parameters were set: The minc (minimum number of genes in a group that will be returned) =15; The parameter summarize we set to “merge”; The results of the DEG patterns for *G. mellonella* larvae RNA-Seq reads mapped to host genome (**Supplementary Figure S10**); and the results of the DEG patterns for *G. mellonella* larvae RNA-Seq reads mapped to pathogen genome (**Supplementary Figure S11**). Similarly, the DEG TSS patterns were analysed for *G. mellonella* larvae reads mapped to the host genome (**Supplementary Figure S12**). The detected gene pattern of the DEG from RNA-Seq is further summarized and grouped in **Figure 6**.

#### Gene Set enrichment and pathway analysis

Gene set enrichment analysis for the host and pathogen was performed using the clusterProfiler R package (V 3.0.4) (7). The function gseGO of clusterProfiler requires an input of the ordered rank gene list from edgeR ordered by logFC. The DEG list from edgeR was annotated with the Gene Ontology terms from Uniprot (8). In addition to the annotated gene list, gseGO requires the AnnotationDbi orgDB package. Since the orgDB was for the *G. mellonella* and *M. mycetomatis* in R Bioconductor, we created two new orgDB for each species. We ran gseGO for each of the gene ontology categories (CC, BP and MF). The number of permutations was specified as 1,000, and the p-value cutoff was defined as “=0.05”. For each of the contrasting time points, the result of the functional enrichment analysis was visualized using dotplot and emapplot. P adjusted value for each plot was shown as well.

#### Method for the identification and analysis of 1,8-Dihydroxynaphthalene (DHN)-melanin, pyomelanin and siderophore biosynthesis pathways in M. mycetomatis

Iron regulation in *G. mellonella* in three biosynthesis pathways in the *M. mycetomatis* were analysed. The pathways are the 1,8-Dihydroxynaphthalene (DHN)-melanin, pyomelanin and siderophore.

Since these pathways are not yet characterized in *M*. *mycetomatis*, we used the fungal *Aspergillus fumigatus* [NCBI: taxid746128] as surrogate species (proxy) to identify the genes involved in the regulation of these pathways. A. fumigatus genes involved in DHN melanin (6 genes) and pyomelanin (6 genes) were fetched from T. Heinekamp and colleagues (9). And for the siderophore biosynthesis pathway, the eight genes fetched from M. Gründlinger and colleagues (10). After collecting the set of genes involved in each biosynthesis pathway in *A. fumigatus*, we looked at their homologous genes in *M. mycetomatis*. Firstly, the protein sequence of *A. fumigatus* genes was retrieved from the UniProt database (8). To obtain the homologous genes in *M. mycetomatis,* we used the NCBI protein-protein BLAST (blastp) web service (11). Blastp run using the following parameters for the chosen search set: The database was non-redundant protein sequences(nr), and the organism used *M. mycetomatis* (NCBI: taxid:100816). After obtaining *A. fumigatus* homologous in homologous genes in *M. mycetomatis* from BLAST, we used the protein-protein interaction database STRING to identify possible functional interaction between the genes for each pathway (12). String stores and provides known and predicted protein-protein interactions (PPI) for thousands of organisms. The protein-protein interactions for homologous genes in *M*. *mycetomatis* enabled us to infer other genes that are interacting with the homologous genes in the PPI. In the PPI, we used the minimum required interaction score = low confidence (0.150) and kept the default value for the rest of the parameters. For each PPI, we obtained the PPI enrichment p-value, the functional enrichments in the PPI (Gene Ontology terms and KEGG pathways), and network characteristics (hub genes, number of nodes, number of edges). Finally, we analysed and visualized the gene expression of homologous genes in *M. mycetomatis* in the list of the differentially expressed genes from RNA-Seq.

#### Validation of selected *G. mellonella* and *M. mycetomatis* genes expression quantitation by Real-Time Quantitative Reverse Transcription PCR (RT-qPCR)

Validation by RT-qPCR of the expression quantitation of selected set of differentially expressed genes of the *G. mellonella* and *M. mycetomatis* was performed. The cDNA was synthesized from 1μg of total RNA with the oligo (dT)20 primer and SuperScript III Reverse Transcriptase (ThermoFisher) following the manufacturer's instructions. RT-qPCR was performed using synthetic primers (**Supplementary Table S4**), TaKaRa Ex Taq HS (Takara Bio), and SYBR Green I Nucleic Acid Gel Stain (ThermoFisher). For each gene in (**Supplementary Table S4**) a DNA fasta file were retrieved from NCBI nucleotide database. The DNA fasta file, used as input for Primer3Plus (13) to pick primers from a DNA sequence. The selected primers by Primer3Plus further processed with Multiple Primer Analyzer (ThermoFisher) and with The Sequence Manipulation Suite (14). PCR was performed for 35 cycles under the following conditions: 98°C for 10 s, 56°C or 64°C for 30 s, and 72°C for 60 s.

### **Supplementary figures**

#### Supplementary figure S1.

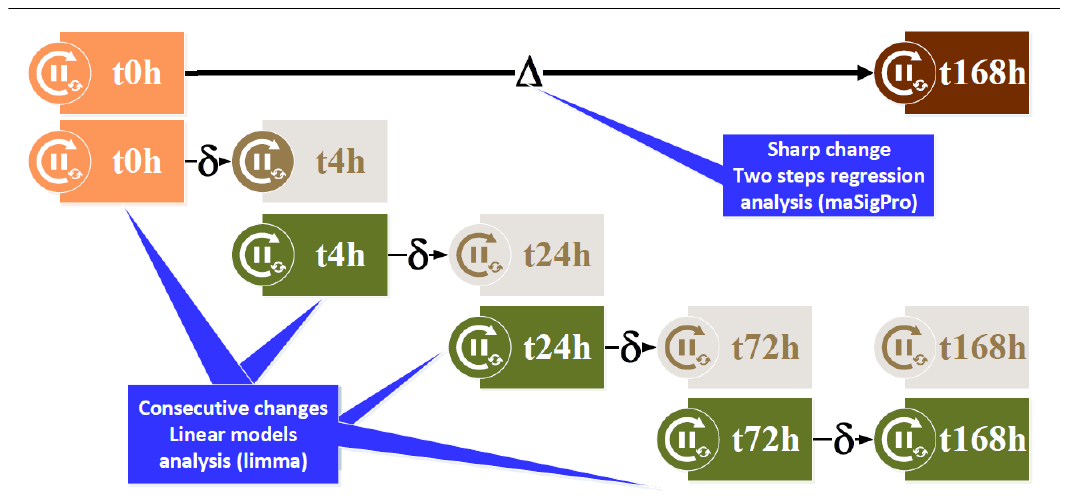

Differential expressing analysis workflow. Combine pairwise comparison with time-course analysis

**pairwise comparison (LIMMA) sharp changes** between time points (Consecutive time points) between every time point and 0 h). T**ime-course analysis (maSigPro) Identify changes over time** (contr Vs. Infected)

#### Supplementary figure S2.

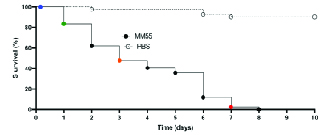

*Galleria mellonella* larval survival curve.

#### Supplementary figure S3.

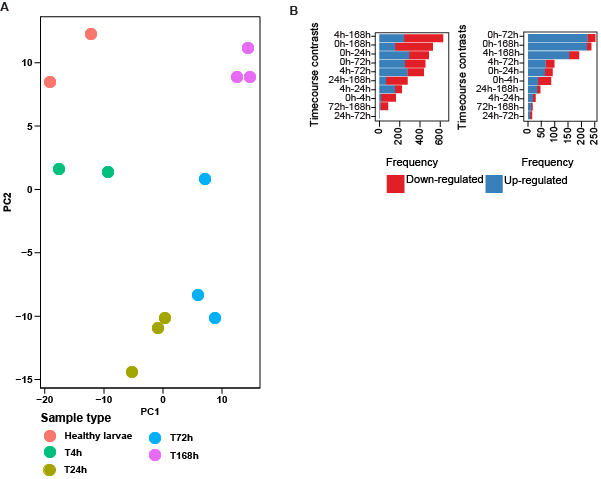

Clustering of LQ-ssCAGE samples and differential expression of the host TSS and enhancers; related to Figure 1 and 2. **A** PCA: LQ-ssCAGE TSSs of host samples; **B** Differentially expressed

host TSS genes (left) and differentially expressed host enhancer genes (right).

#### Supplementary figure S4.

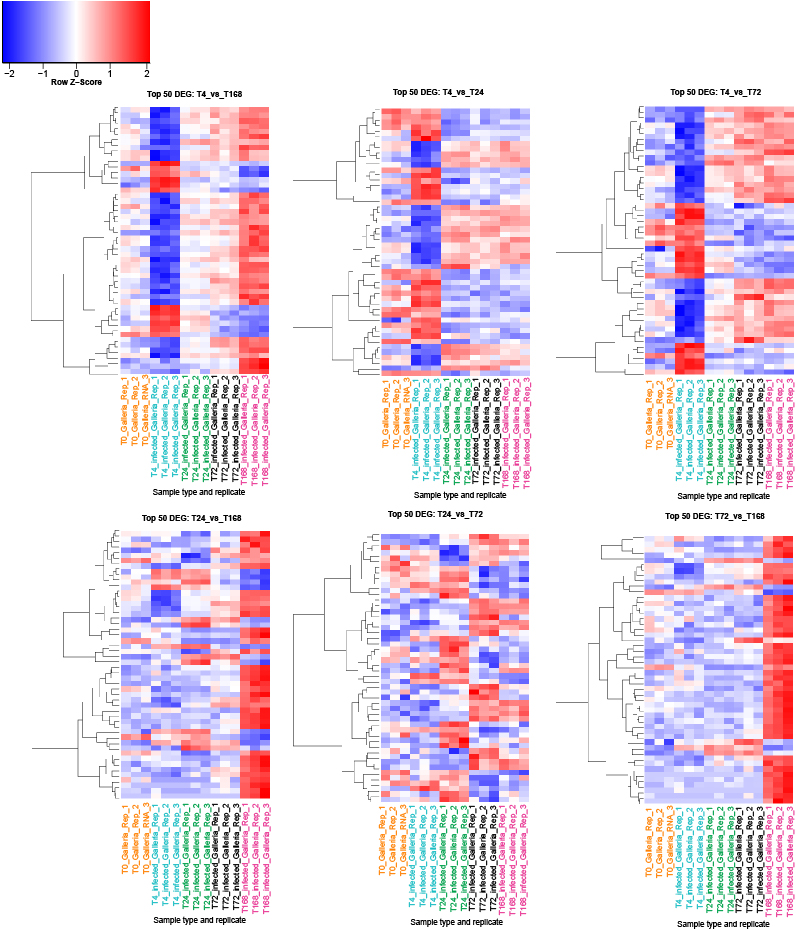

Heatmap of the top 50 host DEG.

#### Supplementary figure S5.

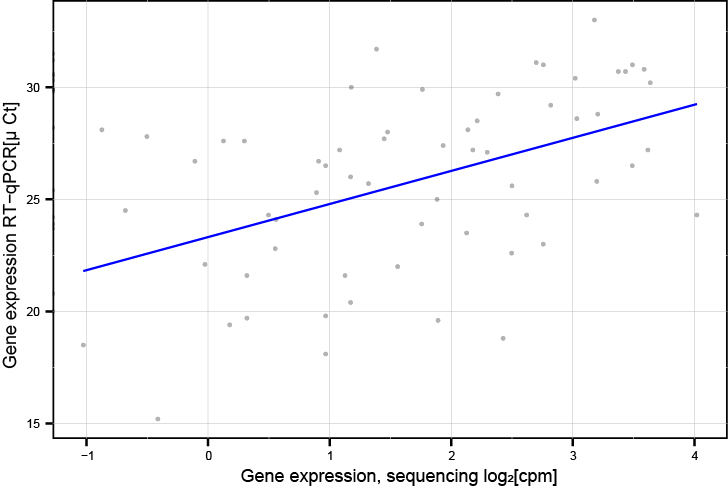

Validation of gene expression quantitation by RT-qPCR.

#### Supplementary figure S6.

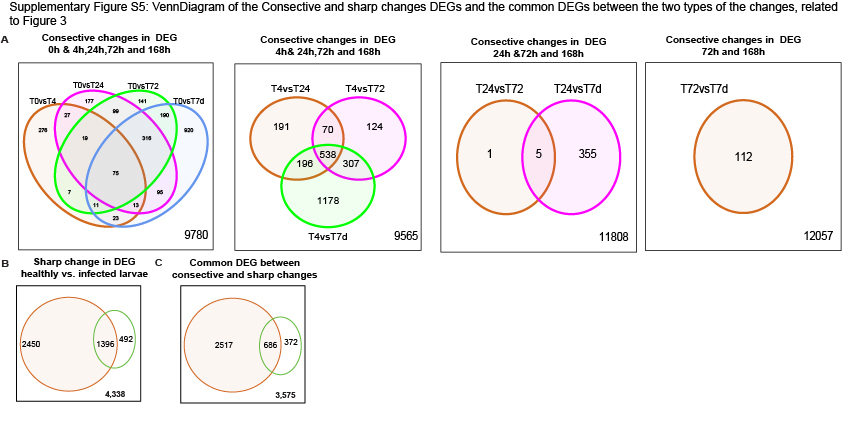

VennDiagram of the Consecutive and sharp changes DEGs and the common DEGs between the two types of the changes, related to Figure 3

#### Supplementary figure S7.

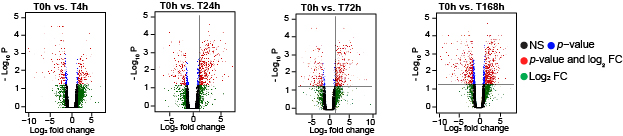

Volcano plot differential expression of the host; related to Figure 3.

#### Supplementary figure S8.

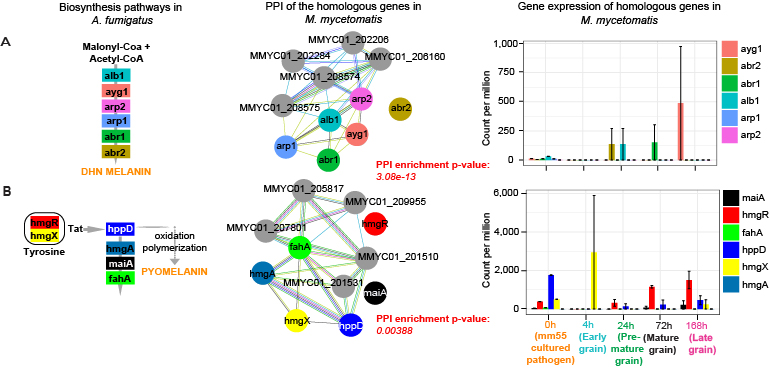

Biosynthesis pathways of 1,8-Dihydroxynaphthalene (DHN)-Melanin and pyomelanin in A. fumigatus with corresponding PPI and gene expression data of *M. mycetomatis* homologues.; related to Figure 7.

#### Supplementary figure S9.

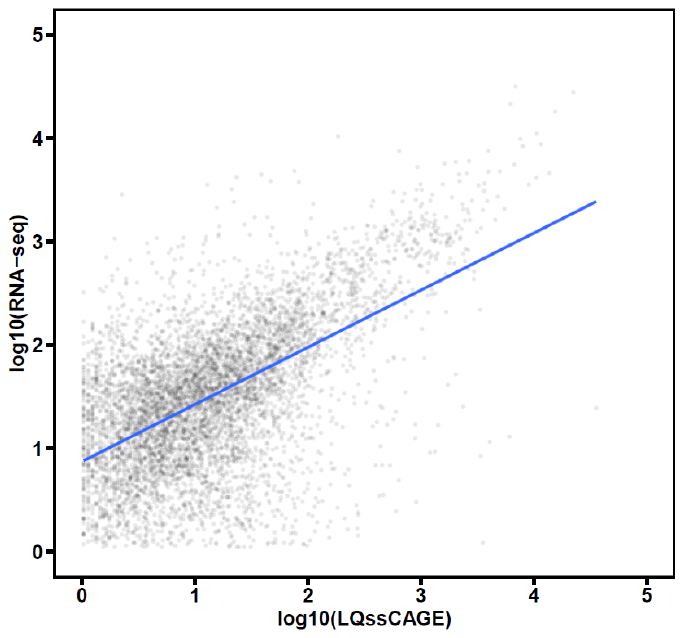

Host reads mapped to GCA_003640425.2 RNA-seq and LQ- ssC A G E correlation; related to Figure 1.

#### Supplementary figure S10.

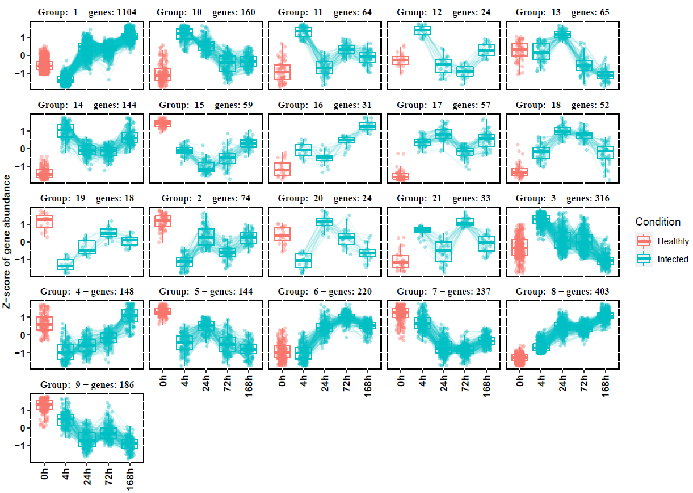

DEG patterns for G. mellonella larvae RNA-Seq reads mapped to host genome.; related to Figure 4.

#### Supplementary figure S11.

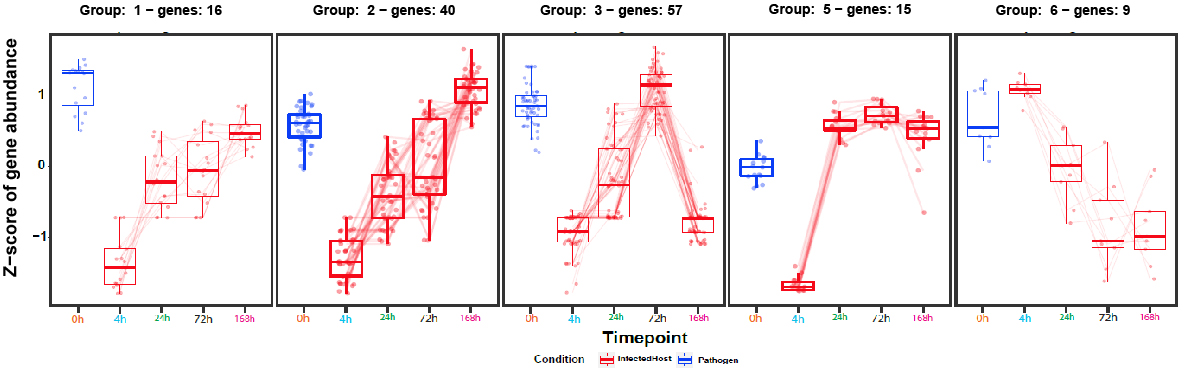

DEG patterns for *G. mellonella* larvae RNA-Seq reads mapped to pathogen genome; related to Figure 4.

#### Supplementary figure S12.

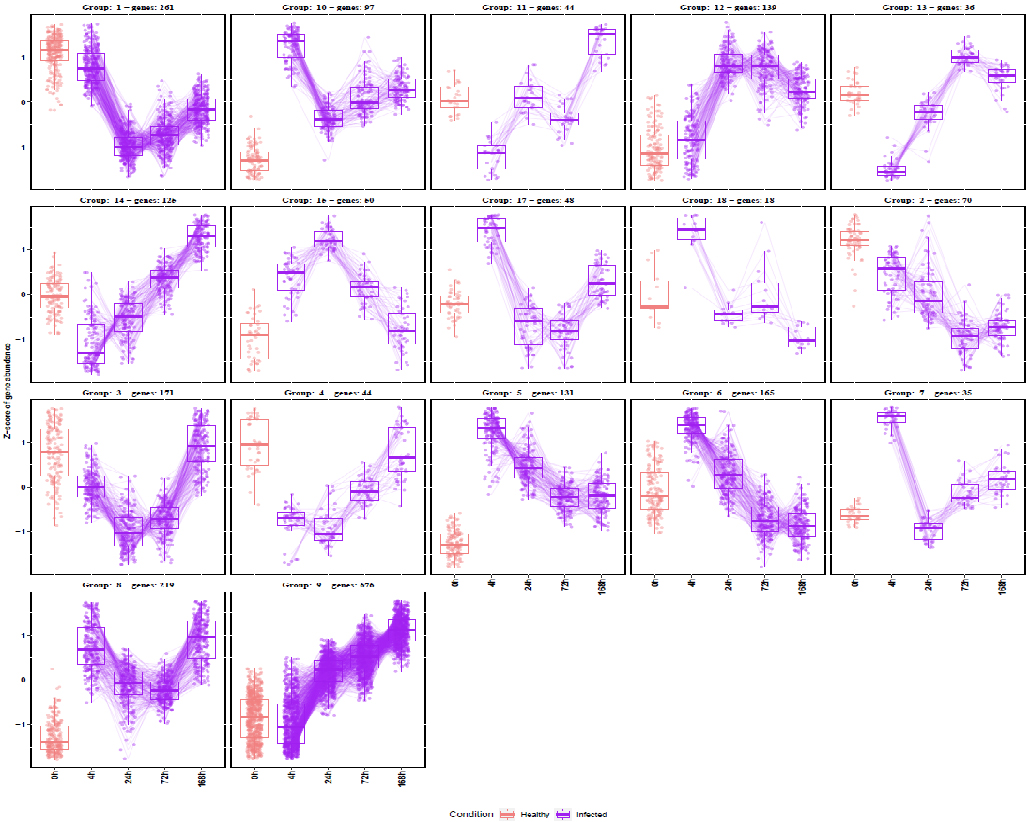

DEG TSS patterns were analyzed for *G. mellonella* larvae reads mapped to host genome.; related to Figure 5.
